# mtDNA-based reconstructions of change in effective population sizes of Holarctic birds do not agree with their reconstructed range sizes based on paleoclimates

**DOI:** 10.1101/2019.12.13.870410

**Authors:** Eleanor F. Miller, Rhys E. Green, Andrew Balmford, Robert Beyer, Marius Somveille, Michela Leonardi, William Amos, Andrea Manica

## Abstract

During the Quaternary, large climate oscillations had profound impacts on the distribution, demography and diversity of species globally. Birds offer a special opportunity for studying these impacts because surveys of geographical distributions, publicly-available genetic sequence data, and the existence of species with adaptations to life in structurally different habitats, permit large-scale comparative analyses. We use Bayesian Skyline Plot (BSP) analysis of mitochondrial DNA to reconstruct profiles depicting how effective population size (*N*_e_) may have changed over time, focussing on variation in the effect of the last deglaciation among 102 Holarctic species. Only 3 species showed a decline in *N*_e_ since the Last Glacial Maximum (LGM) and 7 showed no sizeable change, whilst 92 profiles revealed an increase in *N*_e_. Using bioclimatic Species Distribution Models (SDMs), we also estimated changes in species potential range extent since the LGM. Whilst most modelled ranges also increased, we found no correlation across species between the magnitude of change in range size and change in *N*_e_. The lack of correlation between SDM and BSP reconstructions could not be reconciled even when range shifts were considered. We suggest the lack of agreement between these measures might be linked to changes in population densities which can be independent of range changes. We caution that interpreting either SDM or BSPs independently is problematic and potentially misleading. Additionally, we found that *N*_e_ of wetland species tended to increase later than species from terrestrial habitats, possibly reflecting a delayed increase in the extent of this habitat type after the LGM.

## Introduction

The Quaternary period has been characterised by extensive cycles of glaciation and de-glaciation. The legacy of these ancient large-scale climate alterations is evident today in everything from species’ genetic diversity to population structure (Twitchett 2006; Svenning et al. 2015). Despite the profound impact that past climate changes have had on both flora and fauna, there is limited quantitative evidence on which factors determined how different species fared during these cycles, or how species responded to subsequent post-glacial climate amelioration.

One of the most widely used genetic methods for inferring demographic history is the so-called skyline plot, a family of graphical, non-parametric methods first introduced by Pybus et al. (2000). Grounded in the principles of Kingman’s coalescent theory (Kingman 1982), the ‘skyline framework’ aims to use DNA sequence data to reconstruct a gene tree. The rate of coalescent events within the gene tree can then be used to infer how the population changed in size over time: in essence, periods of low coalescent rates imply a large population while a high density of coalescent events implies a small population. Although skyline plots have been used to reconstruct demographic histories for many species, both extant and extinct (Stiller et al. 2010), and across taxa that include vertebrates (Lu et al. 2012; Vignaud et al. 2014), invertebrates (Sanchez et al. 2016; Villalta et al. 2018) and even bacteria (Segawa et al. 2018), comparative studies across many species are only now emerging (Burbrink et al. 2016).

Skyline plots have been used extensively to infer the response of species during the Last Glacial Maximum (LGM), and they are often paired with climatic reconstructions to infer the changes in available habitat for a given species (Lorenzen et al. 2012; Foote et al. 2013; Calderón et al. 2016). One popular approach for reconstructing possible changes in available habitat for a species through time is the use of bioclimatic Species Distribution Models (SDMs) (Elith and Leathwick 2009). Modelling algorithms combine data on occurrences with environmental data to describe a species’ natural distribution and then simulate how changes in predictor factors may have influenced this available niche space over a period of interest. The underlying logic in linking these approaches is that, assuming limited population structure and appropriate sampling, a skyline plot could, in principle, provide an indication of changes in total population size, and thus of the range occupied by a species. However, the association between effective population sizes (*N*_e_) as reconstructed by skyline plots and species ranges is generally assumed rather than tested.

There are a number of reasons why reconstructed *N*_e_ might not be a good proxy for species ranges. Much attention has been devoted to population structure as a confounding effect, and the recommendation to counter its effects is to pool samples from multiple locations (Heller et al. 2013). However, even with this sampling scheme, there might be a mismatch in the two quantities if mean population density, and thus *N*_e_, was affected by climate change differently to total range extent. For example an increase in mean population density, and thus population size, might occur without a change in range, if the quality of habitat and its carrying capacity increased without a change in its extent (Fig. 1A) (Fordham et al. 2012). Given the positive relationship generally observed between range extent and mean local population density (Connor et al. 2000), *N*_e_ would also be expected to increase by a greater proportion than range extent under climatic amelioration. Another plausible cause of discordance between changes in *N*_e_ and range size is that, without substantial gene flow, skyline plots will mostly reconstruct the population dynamics of the sampled locations rather than the whole species (Miller et al. 2018). Pooling samples from multiple locations can help, but it will not fully resolve the problem (Heller et al. 2013). A plausible scenario that might lead to a disconnect between local *N*_e_ and range size arises during range shifts, as sampled locations, which are suitable for a species at present day, might have been only marginally suitable in the past. In other words, what we now think of as the core area occupied by a species (i.e. where it is abundant, and sampling is more likely) might be inhabited by populations that in the past were at low densities because the local habitat was only marginally suitable. A skyline from such populations would reveal a strong increase in *N*_e_ which reflects the local amelioration of conditions for that species, irrespective of broader range changes (Fig. 1B). A similarly confounded signal will be found in the more extreme scenario where, as a result of a sizeable range shift, the sampled populations inhabit areas which were completely unsuitable in the past, and thus have undergone a founder event after the Last Glacial Maximum (LGM). Such populations would be characterised by a steep increase in *N*_e_ as they recovered from the local bottleneck associated with the founder event (Fig. 1C).

**Figure 1.**
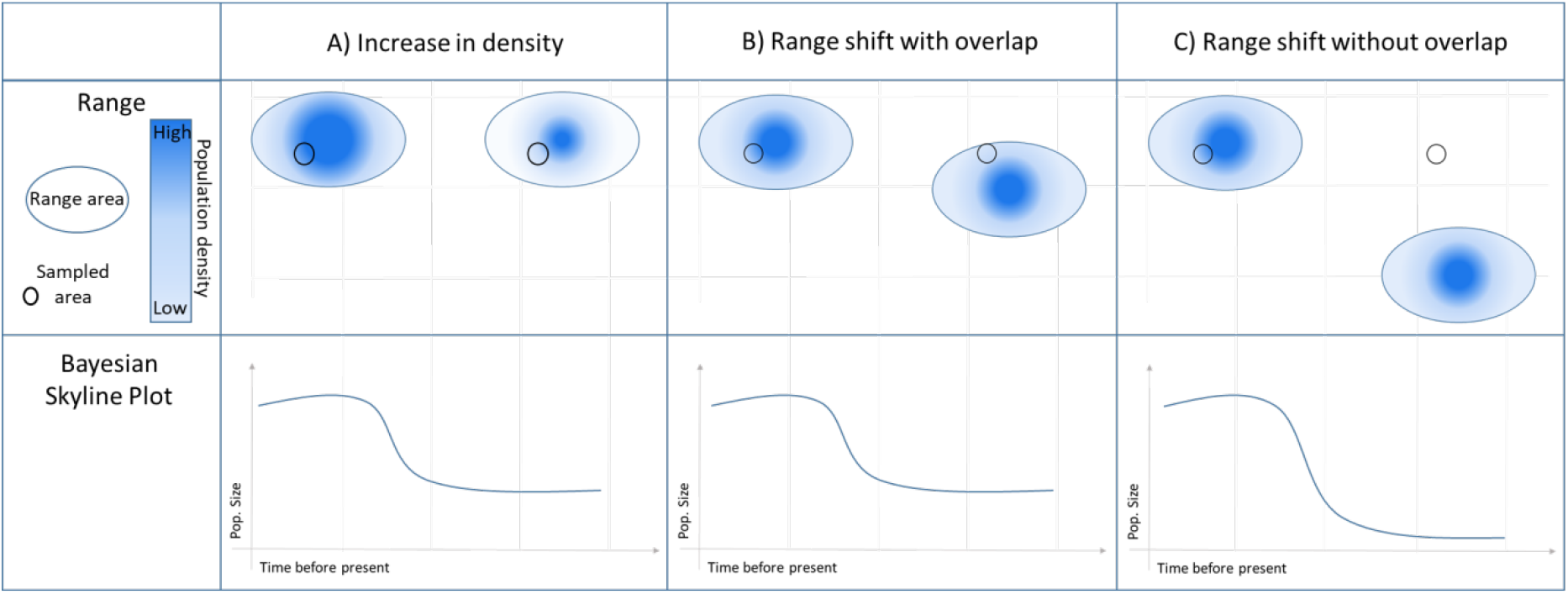
Three scenarios that might lead to an increase in *N*_e_ without a change in range size. The top half of each panel represents a pair of schematic maps of the species range at the LGM (right) and present day (left). The density of colour within each range ellipse shows population density. Circles on the map represent the genetic sampling location. The scenarios are: A) An increase in population size without a change in range recovers an increasing BSP profile. B) Core area today was only marginally suitable in the past, range size remains the same but local amelioration has led to a strong local increase in *N*_e_. C) Sampled populations inhabit areas outside the species range in the past. BSP recovers a steep increase in *N*_e_ associated with founder event.

The extent to which changes in population density and bottlenecks related to range shifts can override the signals linked to changes in range size and the overall metapopulation size is unknown. In the current paper, we mined GenBank to compile a comprehensive dataset of publicly-available mtDNA sequence data from many species of Holarctic birds, and reconstruct their population dynamics using Bayesian skyline plots (BSP) in BEAST2 (Bouckaert et al. 2014). A simple prediction, based on the relative changes of habitat types as reconstructed from the pollen record (Allen et al. 2010), is that species associated with forests (close habitats) should have increased since the LGM, whilst species from open and semi-closed habitats (such as grasslands and steppes) should show a decrease. However, species have more complex niche requirements than a simple association with a broad habitat type, and a more realistic prediction is that *N*_e_ should change in line with changes in extent of the potential geographical range, such as that reconstructed by bioclimatic SDMs. We therefore reconstructed changes in modelled potential range extent between the LGM and the present, using paleoclimate reconstructions and Species Distribution Models, and investigated the relationship between *N*_e_ changes and range size changes. In the case of wetland species, for which reconstructing detailed individual ranges in the past is challenging because they depend upon habitats whose extent is not easily reconstructed, we compared their demographic reconstructions to species from other major biomes to investigate whether there was any consistent pattern in their response to climatic amelioration in the Holocene.

## Results

### Summary of available BSPs

Based on criteria of having a minimum of 10 individuals sequenced for either the NADH dehydrogenase subunit 2 (ND2) or cytochrome b (cytb) genes from the mitochondrial genome (mtDNA), a scan of GenBank yielded a preliminary dataset of 208 species. From these, datasets were discarded for the following reasons: insufficient haplotypes captured for demographic reconstruction (< five), insufficient sequence length (< 200bp), sequences across studies not from comparable sections of the gene, haplotype frequencies not published, an inappropriate sampling strategy used by the original study (e.g. non-random sampling, localised island populations) or extensive population sub-structure (see Materials and Methods for details on the criteria used to select suitable datasets). Application of these criteria left 167 datasets for BSP analyses. All these datasets were analysed with BEAST using a Bayesian Skyline Plot and, with one exception (King Eider, *Somateria spectabilis*), they converged successfully. We note that in BEAST, we adopted a strategy of resizing the ‘*bGroupSizes*’ parameter (see Materials and Methods), potentially constraining the level of detail recoverable in the profiles but ensuring that a large volume of variable quality datasets could be analysed using the same settings (and thus providing comparable estimates).

Data for both ND2 and cytb were available for 28 species. For 18 species, expansion times and profiles across both genes were consistent, and a single profile was then selected to illustrate the population history. For ten species, the two genes showed discordant demographic histories but, in 7 of these cases, one dataset was of appreciably lower quality (e.g. fewer samples, shorter sequences, inappropriate sampling strategy) and was removed. Three species were rejected because the two genes gave discordant profiles despite both appearing to be of comparable quality. See Supplementary Table 1 for details of datasets dropped.

Further profiles had to be excluded as: profiles were either too deep (limit = >1,000,000 years before the present, *n*=9) or too short (limit = <5,000 years before the present, *n*=4) to be informative for the last deglaciation, or profiles showed patterns of expansion or contraction that predated the time period of interest, Marine Isotope Stage 3 (~60 kya, *n*=18)(Van Meerbeeck et al. 2009). Note that, even among the profiles we accepted, there remains great variation in depth due to the sparser and more stochastic branching patterns at the bases of the trees, which cause many profiles either to truncate or to ‘flatline’. Thus, the oldest population size estimates tend to be approximations both because of the reduced information content that impacts all profiles, and the need to use the points of truncation in short profiles or flatline states as the oldest size. For the two species where multiple lineages were identified (pine grosbeak, *Pinicola enucleator,* and horned lark, *Eremophila alpestris*), two separate BSP analyses were performed and an average of the estimates was taken for downstream analysis.

Applying the above filters left 102 qualifying species BSP profiles for further analysis. These species inhabit a wide range of habitats Closed (*n*=43), Open (*n*=17), Semi-closed (*n*=25), Wetlands (*n*=12), and Other (*n*=5); see Materials and Methods for a description of how habitats were grouped. There was no indication that species associated with particular habitats were more or less likely to be excluded (*p* = 0.77, Fisher’s Exact Test, excluding the ‘Other’ category as it had too few species for testing). Skyline profiles encompass a wide range of shapes, variously exhibiting a single sharp point of inflection, gradual changes in size and multiple points of change. No significant differences were found in the proportion of ND2/cytb genes in each habitat type (*p* = 0.78, Fisher’s Exact Test, excluding the ‘Other’ category), nor in the proportions of species from the Palearctic, Nearctic or Holarctic in each habitat type (*p* = 0.10, Fisher’s Exact Test, excluding the ‘Other’ category).

### Direction and magnitude of demographic change

Only 3 out of 102 species showed an overall decrease in *N*_e_ over time, with 7 showing no sizeable change, all other species (*n*=92) increasing to some degree. The direction of change was not associated with habitat (*p* = 0.457, Fisher’s Exact test, excluding the ‘Other’ category), nor was its magnitude (gls *p* ≥ 0. 402 for backbone E and *p* ≥ 0.386 for backbone H, lm without phylogenetic correction *p* = 0.667; Fig. 2A). This result is rather extreme, but it could be the consequence of most species for which several samples are available in GenBank being relatively common and thus having thrived in the Holocene.

**Figure 2.**
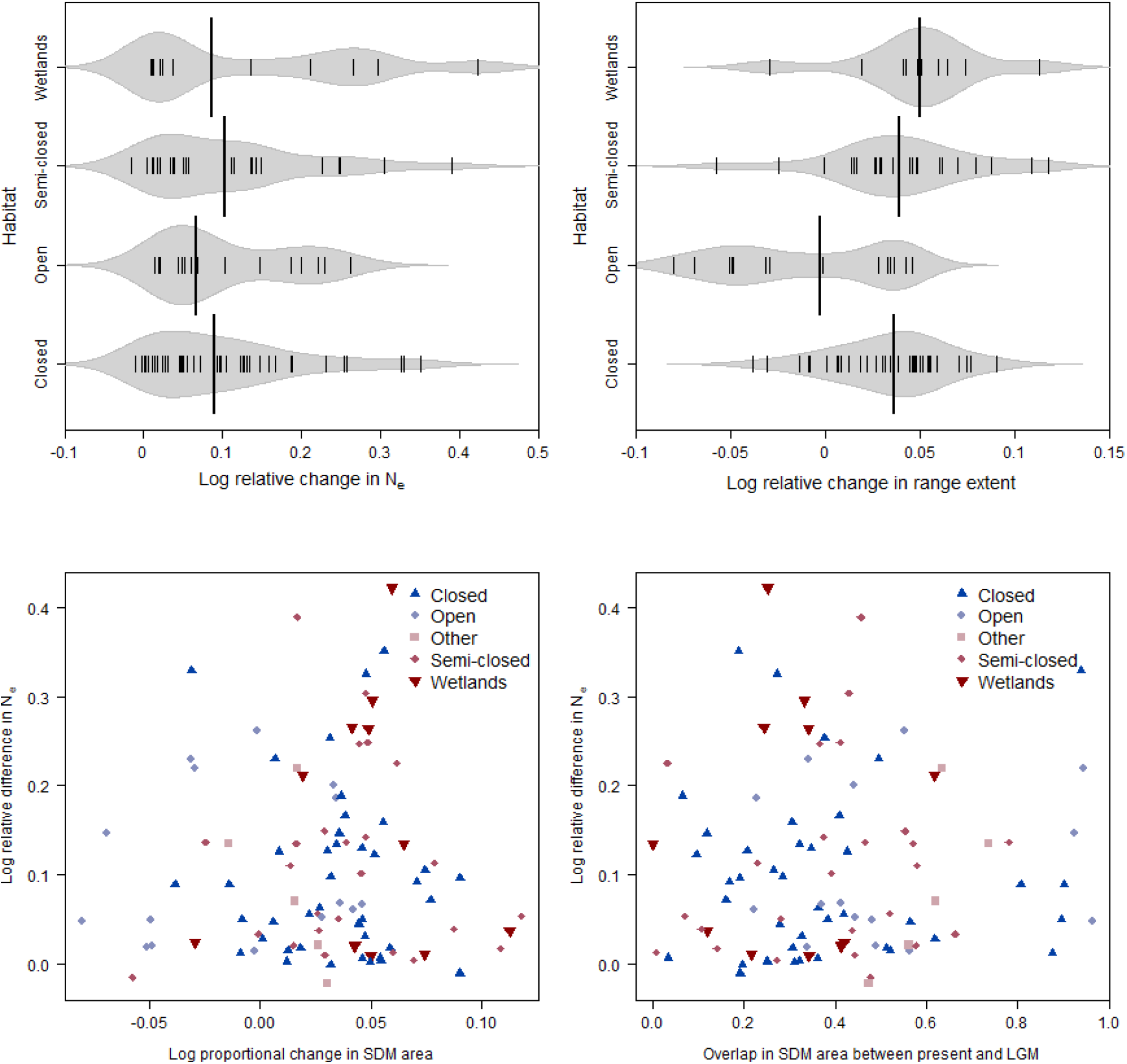
A) Beanplot showing the log relative difference in effective population size (*N*_e_) from 60kya or start of the profile for species from each habitat type. Kernels represent density (i.e. frequency distribution), each small line an individual population, thick black line is median of species-specific values for the given habitat class. Numbers of species per group are; Closed (*n*=43), Open (*n*=17), Semi-closed (*n*=25), Wetlands (*n*=12). B) Beanplot showing the log relative difference in modelled range extent from 21kya for species from each habitat type. Kernels represent density (i.e. frequency distribution), each small line an individual population, thick black line is median of species-specific values for the given habitat class. Numbers of species per group are; Closed (*n*=40), Open (*n*=15), Semi-closed (*n*=25), Wetlands (*n*=11). C) Scatterplot of log ratio of *N*_e_ from 60 kya to 5 kya in relation to the log ratio of change in size of climatically suitable area from 21 kya to the present, based upon species’ individual bioclimate SDMs. D) Scatterplot of log change in *N*_e_ in relation to the proportion of the species’ contemporary range that was also suitable during the LGM. In both scatter plots numbers of species per group are; Closed (*n*=40), Open (*n*=15), Other (*n*=5), Semi-closed (*n*=25), Wetlands (*n*=11).

### Direction and magnitude of change in extent of the potential geographical range

To investigate the plausibility of climate-driven changes in the extent of climatically suitable area contributing to the overwhelming majority of profiles showing an increase in *N*_e_, we created individual Species Distribution Models (SDMs) with the R package ‘*biomod2*’ (Thuiller et al. 2019) for the each of our 102 species. We used occurrences from the GBIF dataset, keeping samples with coordinate data accurate to 10km, removing observations outside the breeding range (defined as the area mapped by Birdlife (BirdLife International and Handbook of the Birds of the World 2018) as being occupied by resident and breeding migratory populations). We extracted for each species uncorrelated environmental variables for the present day and the LGM (R.M. Beyer et al. 2019). We thinned the dataset to reduce spatial sorting bias (Hijmans 2012) and randomly sampled 5 sets of pseudoabsences in the same number as presences from outside the breeding and residential areas. We ran models following four different algorithms (Bagchi et al. 2013) and created ensembles (Araújo and New 2007) by validating each by spatial cross-validation (Roberts et al. 2017). In the end, we had credible SDMs for 96 species; 5 species had to be excluded as we there were insufficient observation points left for analysis after data thinning, and one species was rejected as its SDMs led to a present day projection much larger than the observed range (Supplementary table 2). For the valid SDMs, we generated projected ranges for the LGM (21 kya) and present day and quantified the changes in range size.

As was the case for *N*_e_ changes, the majority of species showed an increase in reconstructed range extent since the LGM (76 out of 96). However, the proportion of species showing an increase in range extent was significantly smaller than the proportion with increased *N*_e_ (*p* = 0.004, data subset to the 96 species for which both analyses were available). Whilst there was variation among groups of species associated with different habitats in the magnitude of change in range extent (gls with phylogenetic correction: *p*≤0.0001 for all 1000 resolutions of both backbone E and backbone H, lm without phylogenetic correction *p* = 0.0001, Fig. 2B) and the direction of change in range extent (*p* = 0.009, Fisher’s Exact Test, excluding the ‘Other’ category), there was no significant match between the direction of the trend in BSP and SDM reconstructions (Supplementary Fig. 1, Fisher’s Exact Test *p* = 0.587). Neither was there a significant positive correlation across species between the signed magnitudes of the changes in the two measures (Fig. 2C; gls with phylogenetic correction: *p*≥0.994 for all 1000 resolutions of backbone E and 1000 resolutions of backbone H, lm without phylogenetic correction *p* = 0.994). Furthermore, taking into account changes in the location of the range (i.e. the proportion of overlap between LGM and present range) also failed to explain the changes in *N*_e_ as reconstructed by BSP (Fig. 2D; gls *p*≥ 0.734 for backbone E and *p*≥ 0.734 for backbone H, lm without phylogenetic correction *p* = 0.734).

### Timing of change

We next explored the relationship between the timing of the dominant population size change and habitat type (excluding the Other category, as it was heterogeneous and only had 5 species). The timings of change in size for each population in the four habitat types are presented in Fig. 3A. Major size change events in wetland-associated species tended to be more recent than for species from the other three habitats (gls *p*≤0.044 in 1000 resolutions of backbone E and *p*≤0.044 in 1000 resolutions of backbone H, lm without phylogenetic correction *p* = 0.044). Similar results were obtained when we excluded species which changed less than 10% in *N*_e_ (which might have added noise) (Supplementary Fig. 2). When using molecular evolution rates from Nabholz et al. (Nabholz et al. 2016) ‘Calibration set 4’ the timing of all expansions are generally consistent with a response associated with the Last Glacial Maximum (LGM). However, we note that using the rate from ‘Calibration set 2’ (which includes older nodes than set 4) would recover older expansion dates (data not shown); given that such dates would correspond to periods of high ice coverage they seem less likely.

**Figure 3.**
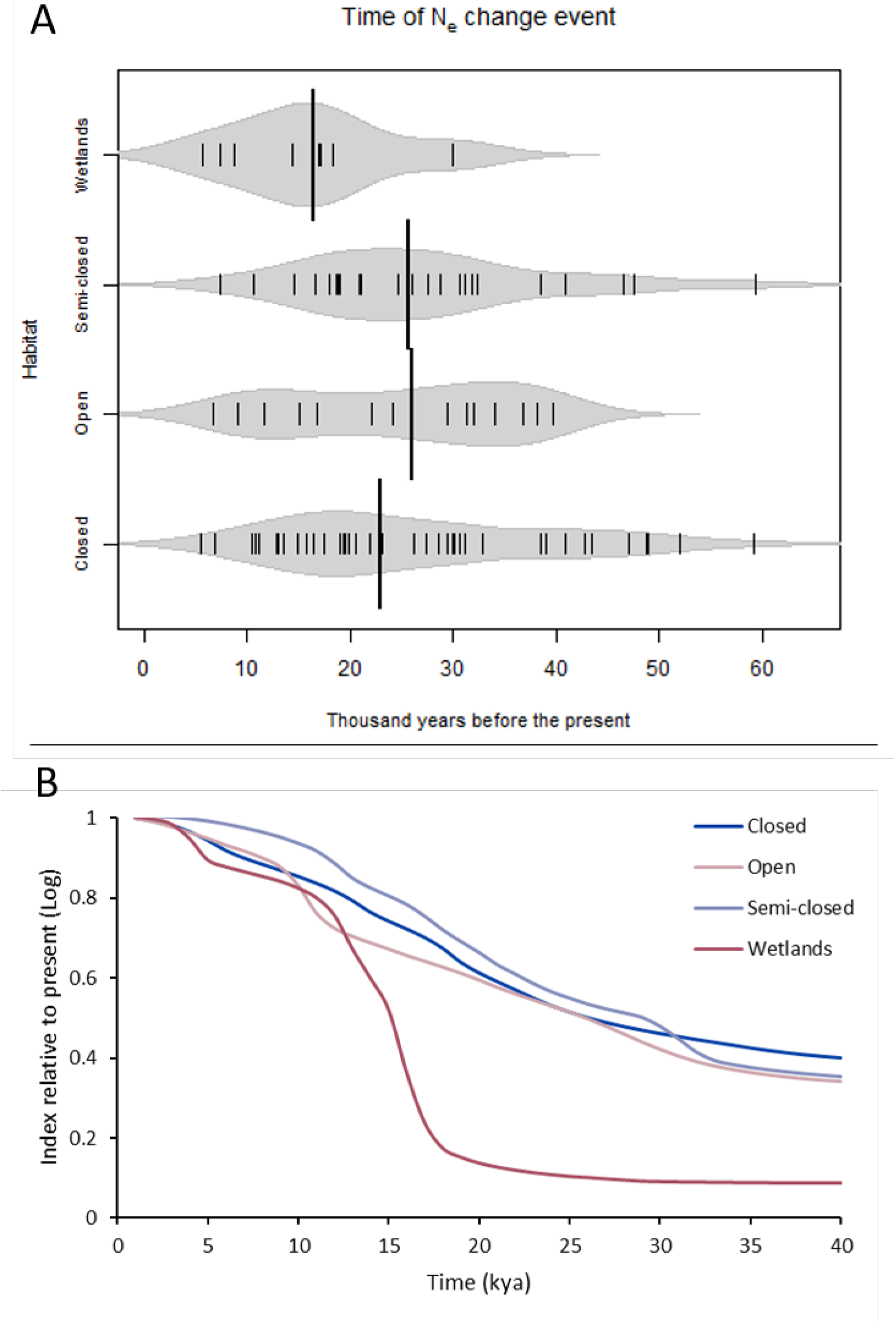
A) Beanplot showing time of dominant effective population size change events for species from each habitat type. Kernels represent density (i.e. frequency distribution), each small line the time of an individual population’s size change event (increase or decrease). Thick black line is median of species-specific change times for a given habitat class. Numbers of species per group are; Closed (*n*=43), Open (*n*=17), Semi-closed (*n*=25), Wetlands (*n*=12). B) A multi-species index (MSI) depicting normalised changed in size averaged across species within each habitat for each time point.

For an alternative view of when the expansions occurred, we further used a multi-species index (MSI). The MSI depicts normalised changes in size averaged across species within each habitat type for each time point (Fig. 3B). Despite exploiting a different aspect of the BSP profile shape, mean change at each time point rather than a single mean date of maximum change, MSI profiles reveal a pattern that is strongly supportive of the previous result where wetland species expand appreciably later than species in the other three habitats.

## Discussion

We generated a large collection of mitochondrial DNA datasets from many bird species to look for evidence of habitat-associated trends in population size through time. Although variable data quality may lead to uncertainties about the magnitude of any particular change in population size we detect, the direction of change is relatively robust (Grant 2015). Out of 102 species, only three species show an overall decrease in effective population size. Changes during the last deglaciation in the modelled extent of the geographical range also indicated increases for most species, though the proportion of increases was lower than for *N*_e_. However, we could find no association across species between the direction or magnitude of change in *N*_e_ and habitat or range reconstructions.

Species with very large ranges at present are the likely “winners” in terms of their response to climatic change and are thus more likely to show an increase in range and population size from the LGM. Widespread species might also be more likely to have been sampled. Indeed, we found that the majority of species we sampled showed an increase in range size based on SDMs. However, the proportion of expanding species according to SDMs was much lower than the one observed for BSPs, thus failing to fully explain the ubiquity of expanding BSPs, and there was no statistical association between changes since the LGM as reconstructed by BSP and SDM.

Colonisation bottlenecks during a range shifts can, in principle, lead to an increase BSP irrespective of the overall change in range size, as long as migration is low enough (i.e. if the BSP captures the local dynamics in a given population/small geographic region rather than the whole range). However, if this mechanism was important, we would expect species exhibiting an increase in BSP despite a range contraction to be associated with large shifts; this is not the case (Supplementary Fig. 3). Therefore, colonisation bottlenecks do not seem to explain the ubiquity of expanding BSPs in our dataset.

Increases in migration can also lead to an increasing BSP without any change in census population sizes. The potential role of migration in producing counterintuitive *N*_e_ estimates when assuming panmictic populations for a whole species has received much attention recently in the context of interpreting cross-coalescence (MSMC) profiles (Mazet et al. 2015). However, it is difficult to envision a scenario where migration would increase significantly in the face of a range contraction; the effect of migration is more likely to be seen during an expansion, when previously isolated fragments are reconnected. Thus, it seems unlikely that migration can explain the ubiquity of increasing BSPs in Holarctic birds.

A final, more likely but difficult to test explanation for our results is that population densities have increased since the LGM. Thus, even for species that have experienced a range contraction, there might have been changes in local population dynamics such that the average density is higher at present. This decoupling of range extent and density makes interpretation of the SDMs and BSPs very challenging. To resolve them, there is a need to use species abundance models that explicitly predict population densities rather than presence/absence (Potts and Elith 2006; Howard et al. 2014; Johnston et al. 2015). Whilst fitting such models is possible in principle, they have been little used because extensive population density information is rarely available for any given species. However, recent efforts such as those by the Cornell Lab of Ornithology (https://ebird.org/science/status-and-trends/)(Auer et al. 2019) have started collating such information, opening a window in better understanding the link between range size and density. Our results, however, do raise a caveat in the interpretation of SMDs and BSPs for extinct species; arguably, the best strategy is to couple the two approaches, as only their combined results might provide a good overview of the fate of a species.

The timings we find for when population expansions occurred agree broadly with those of changes in climate after the LGM. Dates were based on mitochondrial mutation rates calibrated for body size and based on a calibration set that included relatively young species splits (Nabholz et al. 2016), and thus likely to give faster mutation rates (i.e. less affected by selection) that were appropriate for within species analysis (Howell et al. 2003; Penny 2005; Ho 2007). We note that using a calibration set that included older species splits (Nabholz et al. 2016) would lead to much older (well before the last glaciation), and thus less realistic changes in effective population sizes. However, we strongly caution the reader that mutation rates calibrated by bird body size, whilst the best available option for comparative analysis, are likely to be very noisy, and individual species estimates should not overly interpreted. Ideally, we would need taxon-specific mutation rates (Hope et al. 2014) which are simply not available for the number of species we are investigating in this study. Having said this, the fact that species from the same habitat tend to yield broadly similar profiles gives us confidence that the relative timings are likely robust, even if the absolute values still have room for improvement.

We found that changes in *N*_e_ for wetland-associated species have occurred more recently than those from terrestrial habitats. Although the significance is marginal when based on point estimates for the date of most rapidly changing size, the finding is supported by multispecies index analysis, which also reveals a pattern of later expansion among wetland species. Compared to many terrestrial habitats, wetlands tend to be less stable. Factors such as local water table levels, the amount of meltwater from retreating ice-sheets and rates of soil erosion all play into wetland habitat development and could have delayed the establishment of stable wetland habitats after the LGM. Although reconstruction of wetland environments and the modelling of wetland recovery is difficult (Kaplan 2002; Valdes et al. 2005; Lafleur 2008; Fan and Miguez-Macho 2011), analysis of pollen across Eurasia shows that species associated with wetlands such as Sphagnum moss and Alder trees both exhibit much later expansions compared with terrestrial species (Allen et al. 1999; Giesecke et al. 2017). Indeed, the expansion of alder relative to other trees (Allen et al. 1999) matches closely the relatively later expansion we find for wetland versus non-wetland birds.

BSP analysis is powerful but depends on a number of assumptions that are rarely met in real data, most notably the use of a random sample of individuals drawn from a panmictic population (Pannell 2003; Chikhi et al. 2010; Heller et al. 2013). Consequently, most profiles should be seen as approximations that are easy to over-interpret (Grant 2015; Miller et al. 2018). However, increasing numbers of public domain datasets open the door for studies based on characteristics averaged across multiple profiles constructed from species or populations that share a common habitat or other trait. This averaging approach is not without its own significant challenges and the data still need stringent filtering. In our study, we constructed and inspected network diagrams for each dataset, allowing species with genetic outliers and evidence of strong population substructure to be identified and either divided or excluded. The large number of species investigated allowed us to see a clear pattern of population expansion in almost all species following the LGM, irrespective of their range dynamics, and a tendency for the expansion to occur later in wetland species. The near-ubiquitous signal of expansion suggests a decoupling of range size and local densities, implying a need for carefully interpretation of BSP to describe species-wide responses.

## Materials and Methods

### Raw Genetic Data

We assembled two databases, one for NADH dehydrogenase subunit 2 (ND2) and one for cytochrome b (cytb); these two genes are among the most frequently uploaded avian mtDNA loci in GenBank. We first collated summary information on all available avian ND2 and cytb sequences in GenBank using a custom R script, and then screened it for Holarctic species using the list of Voous et al. (Voous 1977). We only retained species with more than 10 accession for either gene; when a species had sufficient data for both ND2 and cytb, sequences for each gene were extracted and handled as distinct datasets.

### Alignment

Sequence data for each species / gene combination come from multiple independent studies and often differ in the gene region they analyse. Comparable regions were found by aligning sequences in MEGA (version 7.0; (Kumar et al. 2016) using the programme ClustalW (Thompson et al. 1994). Sequence data for each taxon were then trimmed to the longest common section between all samples. If inclusion of a single sequence required the loss of >200bp from more than 50% of the other sequences, that sample was excluded. Furthermore, all positions containing insertions, deletions or sequencing ambiguities were removed. When studies uploaded only one copy of each haplotype, we used haplotype frequencies from the associated publications to generate the appropriate number of copies in our database. Publications that lacked haplotype frequency were excluded from the analysis. After frequency correction the available sample sizes varied from 11 to 453 sequences per species, with lengths from 236 base pairs (bp) to 1137 bp.

### Median Joining Networks

For each species / gene combination, we built a median joining network (MJNs) in POPART (Leigh and Bryant 2015). If the MJN contained long branches, defined as 30 or more nucleotide substitutions on a single branch, the sampling location for that species was reviewed because such long branches are indicative of profound population substructure. If clear geographical separation or grouping was found, the data were divided as appropriate and treated as discrete datasets. Single samples with >30 mutations on a branch were considered extreme outliers and dropped from alignments.

### Mutation Rate

Recent work (Nabholz et al. 2016) proposes that body-mass can be used to inform more accurate calculations of taxon-specific substitution rates and provides a correction factor for variation in rates according to body mass as well as major mtDNA loci. We created dataset-specific molecular evolution rates using the body mass / gene correction factors from Nabholz et al. (2016) ‘Calibration set 4’ (3^rd^ codon position), as it includes younger species splits that should lead to estimates more appropriate for within the within species dynamics that we investigated in our paper. Due to the uncertainty surrounding mutation rates, analyses based on ‘Calibration set 2’ were also run (data not shown). Body mass data was taken from Dunning et al. (2007).

### BSP Analysis

Whilst there now exist a range of related skyline plot methods (see Ho and Shapiro 2011 for a technical review) we focus on BSPs (Drummond et al. 2005). The relative simplicity of this approach, and its inherent robustness, makes it particularly suitable for the heterogenous quality of datasets investigated in this paper. For each dataset, BSP analyses were implemented in BEAST2 (Heled and Drummond 2008; Bouckaert et al. 2014) using a strict clock with a taxon-specific body mass / gene mutation rates, run lengths of 300 million steps sampled every 30,000 steps, with the first 10% discarded as burn-in. The integrated Bayesian application bModelTest was used to select the most appropriate site model and parameters for individual analyses (Bouckaert 2015) and ‘*bGroupSizes*’ was set to 3. All other parameters were left as default. Each analysis was run twice and convergence verified by both a visual inspection of MCMC trace output in Tracer v1.6 and confirming that the effective sample size (ESS) values exceed 200 (Drummond and Rambaut 2007). Demographic reconstructions were then summarised in Tracer v1.6 (http://tree.bio.ed.ac.uk/software/tracer/) with ‘*Number of bins*’ set to 500, and plotted in R.

### Inclusion Criteria

Where data were available for both ND2 and cytb BSP profiles were compared along with summary statistics on each dataset. When profiles were in agreement, the best supported dataset was retained, e.g. largest sample size, longest sequences, to represent that species’ history. If profiles were not concordant but there was a clear disparity in the quality of the datasets, we again kept the profile from the better dataset. If the profiles did not show similar trends, and there was no difference in the data quality, we conservatively rejected both profiles. For inclusion in further analysis, profiles needed to have a history deeper than 5 thousand years ago (kya) but shallower than 1 million years ago, and also have recovered a change event (increase and / or decrease) dated within the last 60 kya.

### Habitat Classification

The species we considered were associated with a wide range of habitat types, especially in terms of the predominant vegetation. Given the need to use a small number of habitat categories, no classification system can be perfect. We used the expert ornithological opinion of one of us (REG) to classify each species according to the major habitat with which each species is currently associated, based upon descriptions of their natural habitats in a standard work (BirdLife International and Handbook of the Birds of the World 2018). After initial data quality filtering 138 species were available to be classified this way. The habitat classes selected were Closed (forests), Semi-closed (shrubland and open woodlands), Open (grassland, montane and steppe), and Wetlands (freshwater wetlands). Some species could not be placed in one of these classes and there were other classes (e.g. Rivers) with five or fewer species. These exceptions were grouped into a category referred to as ‘Other’.

### Timing of Expansion

Identifying a population’s point of expansion from a BSP profile proved difficult given the wide range of shapes present: some populations changed little or very gradually in size, while others showed sharp and / or multiple points of inflection. We chose to identify inflection points using a custom algorithm, however, as timings could be confounded by the capture of local optima we also used visual inspection of the plotted data and a custom R script to review and extract exact timings. The ‘algorithmic’ method used a moving window approach, repeated for five different window sizes. In each window a linear regression was fitted, and its equation used to remove any slope, before fitting a second order polynomial and recording the difference in y-axis value between the mid-point and the average start and end of each window. This method usually identifies one point of maximum turn (increase or decrease) but the visual inspection of each plot allowed us to capture any additional size change events that were present.

### Size change

To compare the magnitude of estimated population expansions within the period of interest, the relative population size change between 60 kya and 5 kya was calculated. *N*_e_ at 60 kya and 5 kya was interpolated for each BSP profile and where profiles were shorter than 60 kya the population was assumed to be the size at the start of profile. Given the uncertainty of molecular methods we prefered to use size estimates from 60 kya / the earliest possible point instead of estimating the size at the height of the LGM (21 kya) where it is more likely the analysis would catch profiles already undergoing demographic change as a result of climatic events.

We also created a multispecies index (MSI) to further explore the changes in *N*_e_. MSI is a form of average profile based on the average ratio of estimated population size between adjacent time points. Specifically, for each pair of adjacent time points we calculate the geometric mean of the ratios of all species for which data exist. We reconstructed a profile for each broad habitat grouping based on these ratios, working back from the present which is assigned a value of 1.

### Phylogenetic Correction

The species included in our study are not phylogenetically independent, so all analyses included a phylogenetic correction based on the most complete molecular phylogeny of extant birds (www.birdtree.org, (Jetz et al. 2012)). Jetz et al. (2012) present two phylogenetic backbones, the Ericson and Hackett backbones (‘backbone E’ and ‘backbone H’ from now on), and, as we considered both to be equiprobable, all analyses were repeated with both. For each backbone, we generated 1000 trees, randomly resolving polytomies in each, and repeated all analyses for each tree with a *“pgls”* phylogenetic correction from the Caper package in R. For all analyses, we also provide the results based Ordinary Least Squares (i.e. without phylogenetic correction).

### Species Distribution Models

The whole pipeline is provided as a commented R script in the supplementary materials.

#### Present day and LGM paleoclimate reconstructions

In order to identify areas suitable for species through time, high-resolution climate data from the past is needed. We used a 0.5° resolution dataset for 19 bioclimatic variables; Net Primary productivity (NPP), Leaf Area Index (LAI) and all the BioClim variables (Hijmans et al. 2005) with the exclusion of BIO2 and BIO3; covering the last 21,000 years in 1,000 year time steps (R.M. Beyer et al. 2019). This dataset was originally constructed from a combination of HadCM3 climate simulations of the last 120,000 years, high-resolution HadAM3H simulations of the last 21,000 years, and empirical present-day data. The data were downscaled and bias-corrected using the Delta Method (R. Beyer et al. 2019).

#### Species data preparation

Species occurrences were downloaded from the GBIF database (https://www.gbif.org) without any preliminary filtering (download links are available in Supplementary table 2). Occurrences were then filtered based on the accuracy of the coordinates (maximum error: 10 km), keeping only observations within the breeding and resident geographical ranges from Birdlife (BirdLife International). The occurrences were then regridded based on the palaeoclimatic reconstructions (0.5 x 0.5 degrees) and, as the method used works on presence / absence and not frequency, only one presence per grid cell was kept.

For each species, this cleaned dataset of presences was then used to select a subset of bioclimatic variables from the 19 variables available in the paleoclimatic reconstructions (R.M. Beyer et al. 2019). In order to avoid using highly correlated variables, which may increase noise in the data (Guisan et al. 2017), a correlation matrix was constructed between the variables associated with each presence. Where two values were highly correlated, the variable with the lowest overall correlation across the matrix was retained and this way a set of uncorrelated variables (threshold = 0.7) were selected.

Opportunistic observations, such as those collected in the GBIF database, tend to have geographic biases in sampling effort. In order to reduce the risk of geographic sample bias affecting the SDMs, the dataset was thinned using the R package *spThin* (Aiello-Lammens et al. 2015), a minimum distance of 70 km was enforced between observations. Given the random nature of removing nearest-neighbour data points, this process was repeated 100 times (‘rep’ = 100) but in order to keep as much information as possible the result which kept the maximum number of observations after thinning was retained for downstream analysis.

#### Species Distribution Model fitting

Species Distribution Modelling was performed using the R package *biomod2* (Thuiller et al. 2019) for all species with more than 10 occurrences after filtering and thinning (Stockwell and Peterson 2002). The thinned dataset was used as presences, the whole region (i.e. the land mass of Eurasia or North America) as background, and then the same number of pseudo-absences as presences were randomly drawn of from outside the BirdLife resident and breeding masks 5 times, creating 5 independent datasets for further analysis. We found that the most effective strategy to retrieve SDMs for the present day that were consistent with the best available estimates of species’ modern day range, as provided by BirdLife resident and breeding masks, was to draw pseudo-absences from outside these masks. By confirming that the estimated distributions recovered for the modern day were in accord with the BirdLife range predictions we were confident that the modelled niche being projected to the past was with as accurate as possible (an example using the *Passer domesticus* dataset is given in Supplementary Fig. 4).

Following Bagchi et al. (Bagchi et al. 2013), models were run independently for each of the five pseudoabsence datasets using four different algorithms: generalised linear models (GLM), generalized boosting method (GBM, in the mentioned reference defined as “boosted regression tree”), generalised additive models (GAM) and random forest. Model evaluation was performed by spatial cross validation (Roberts et al. 2017), i.e. splitting the dataset (both presences and all five runs of pseudoabsences) based on latitudinal bands in America and longitudinal bands in Eurasia (Supplementary Fig. 5) with the R package *BlockCV* (Valavi et al. 2019), and using 4/5 of the splits to calibrate the model and the remaining 1/5 to evaluate it. A data split cannot be used for evaluation if it contains only absences. For this reason, given the great variety of distribution of the species analysed, we decided to maximise the probability of having at least some presences in all data splits by creating 15 spatial blocks encompassing the whole region of interest, either North America (East-West bands) or Europe (North-South bands). Each block was given an ID, numbered sequentially 1-5, the 15 blocks were then grouped into five working data splits by these ID number.

The models were run based on each of the four mentioned algorithms five times (once for each pseudoabsence run), using in turn four of the five defined data splits to calibrate and one to evaluate based on TSS (threshold = 0.7).

A full ensemble, combining all pseudoabsences sets and algorithms (Araújo and New 2007), was then built using only models with TSS > 0.7 averaged through four different statistics: mean, median, committee average and weighted mean. The statistic showing the highest TSS was then projected to either Eurasia or Northern America considering both the present-day climate and the palaeoclimatic reconstruction for the LGM (defined as 21 kya).

#### Range change and overlap

Finally, the binary projection was used to estimate the climatically suitable area (in square kilometres) for each species both now and in the LGM. In order to do so, we first re-projected the rasters to the Eckert IV equal-area pseudocylindrical projection setting the grid size to 50×50 km, and then multiplied the number of cells occupied in each period, and their overlap, by the cell area (2500 km^2^).

## Supporting information

Pipeline for SDM analyses

## Data Availability

All genetic sequence data used in this study were downloaded from the GenBank website — https://www.ncbi.nlm.nih.gov/genbank/

All species occurrence data were downloaded from the Global Biodiversity Information Facility (GBIF) database; DOIs are available for each individual species dataset in the Supplementary file ‘BSP_and_SDM_info.csv’ (Supplementary Table 2).

## Author contributions

E.F.M. performed the analyses, produced the plots and wrote the manuscript. All SDMs were constructed by M.L. Expert ornithological knowledge was provided by R.E.G. who also proposed the MSI analysis. All authors provided supervision and comments on the manuscript.

## Acknowledgements

We thank Professor Brian Huntley and a group of paleoecologists gathered to honour him upon his retirement for pointing out the similarity of our finding of a later increase in effective population size of wetland birds and a similar delay in the expansion of some wetland plants during the last deglaciation.

## Additional information

The authors declare no competing financial interests.

