## Supplementary figures and table 1 for "mtDNA-based reconstructions of change in effective population sizes of Holarctic birds do not agree with their reconstructed range sizes based on paleoclimates"

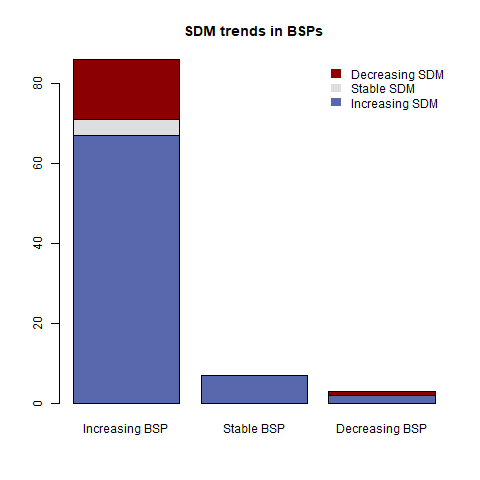

Supplementary Figure 1. Barplot of BSP trends coloured by proportions of plots that have an increasing, stable or decreasing SDM trends. It was not possible to construct SDMs for all species, therefore *n*=96.

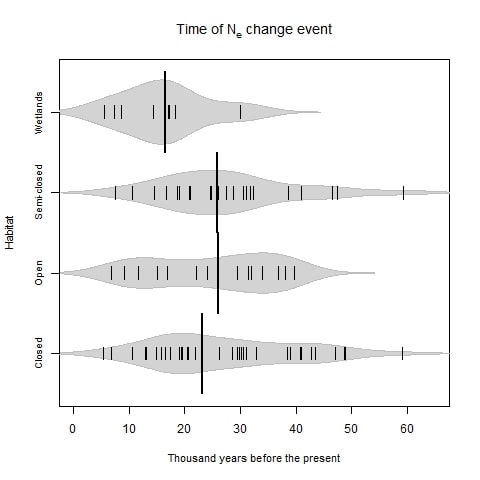

Supplementary Figure 2. Beanplot showing time of population change event for species from each habitat type, excluding populations with an overall size change less than 10%. Kernels represent density, each small line the time of an individual population’s size change event (increase or decrease). Thick black line is median time for change in the habitat. Numbers of species per group are; Closed (*n=*37), Open (*n=*17), Semi-closed (*n=*24), Wetlands (*n=*12).

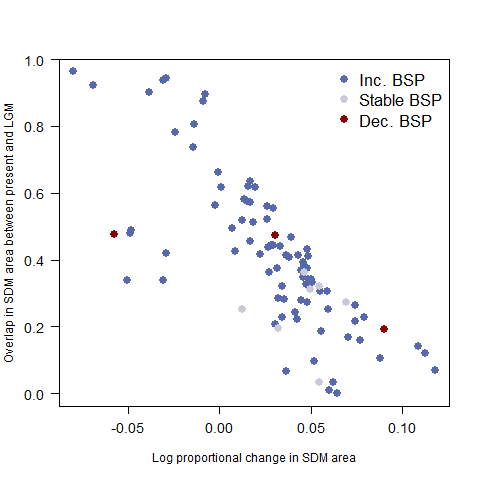

Supplementary Figure 3. Scatter plot of change in overall SDM area and the proportion of each SDM in present that was also suitable for that species at the LGM (21 kya). Points are coloured by trend in *N*_e_ from BSP.

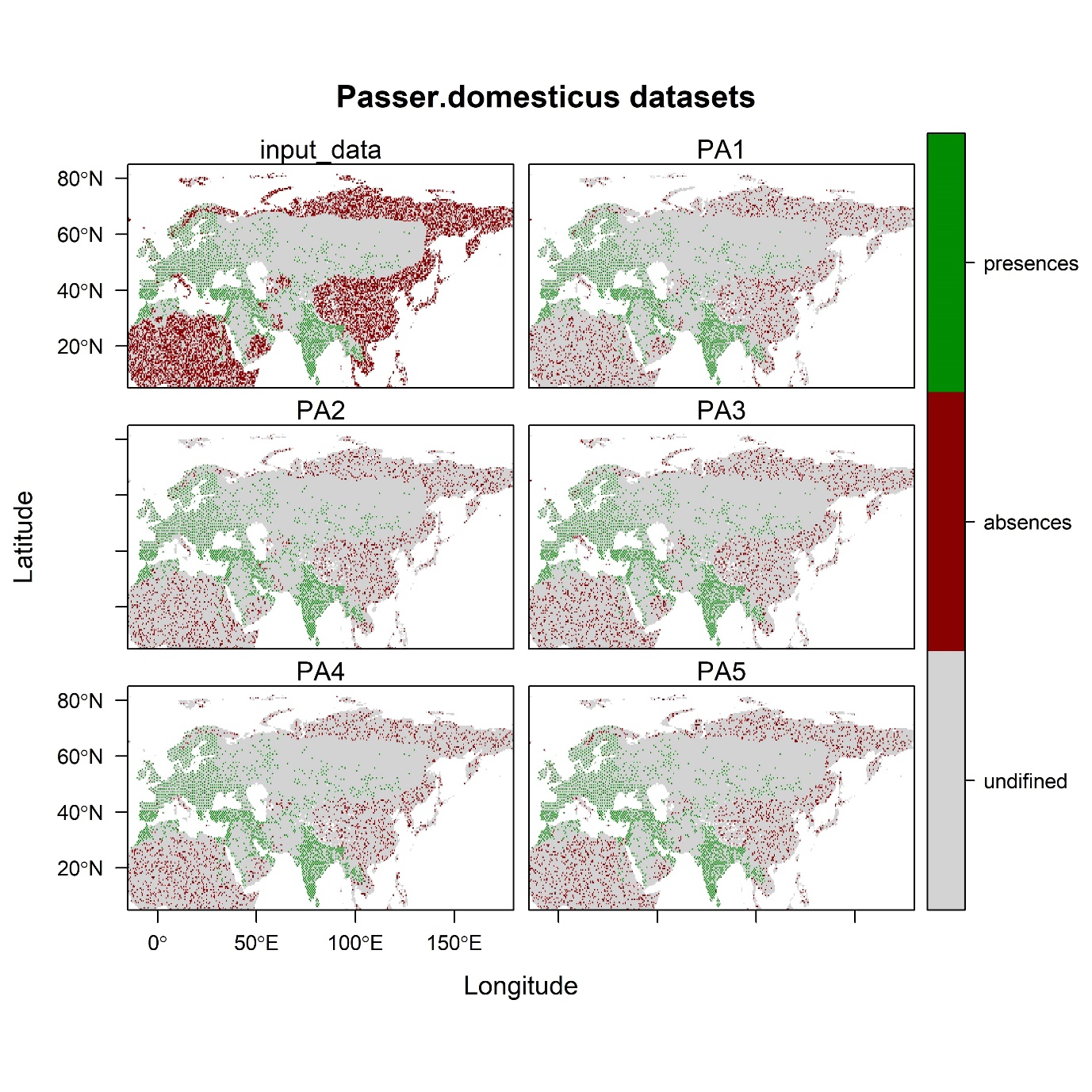
Supplementary Figure 4. An example of the BirdLife resident and breeding masks, species shown is the house sparrow (*Passer domesticus*). Panel one shows all data, PA1-5 show different sets of randomly sampled pseudoabsences.

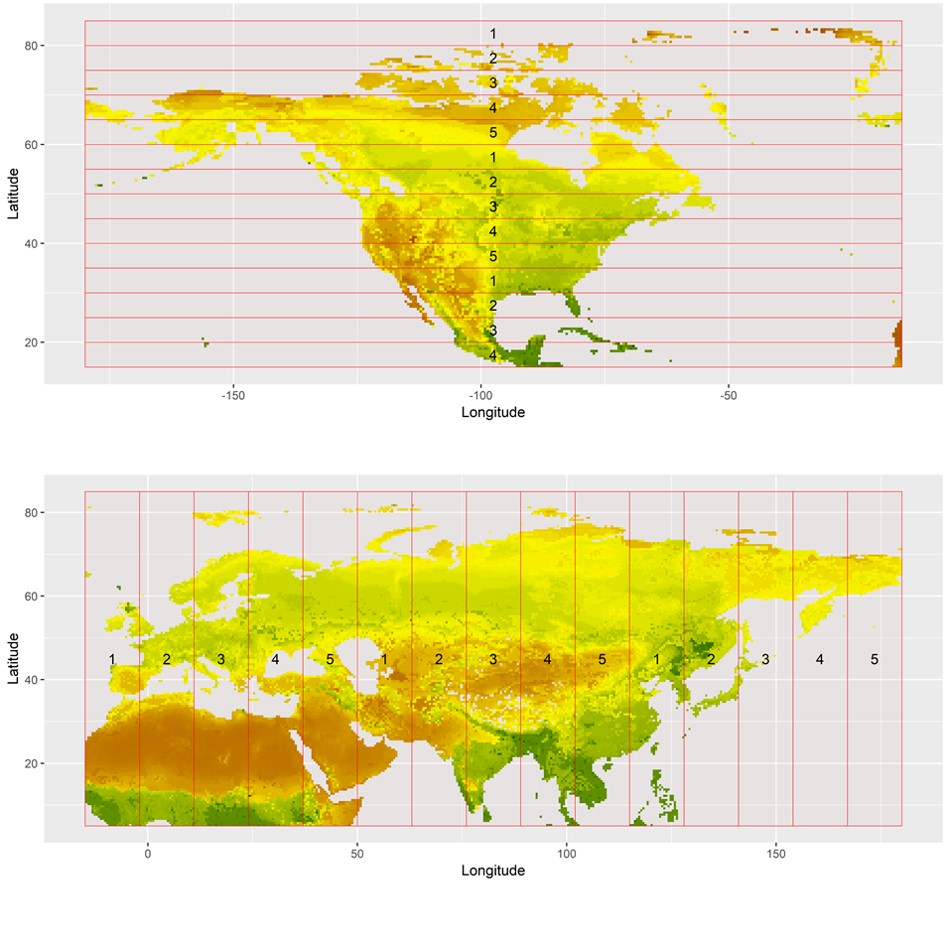

Supplementary Figure 5. Spatial blocks based on latitudinal bands in America and longitudinal bands in built with the R package *BlockCV* (Valavi et al. 2019).

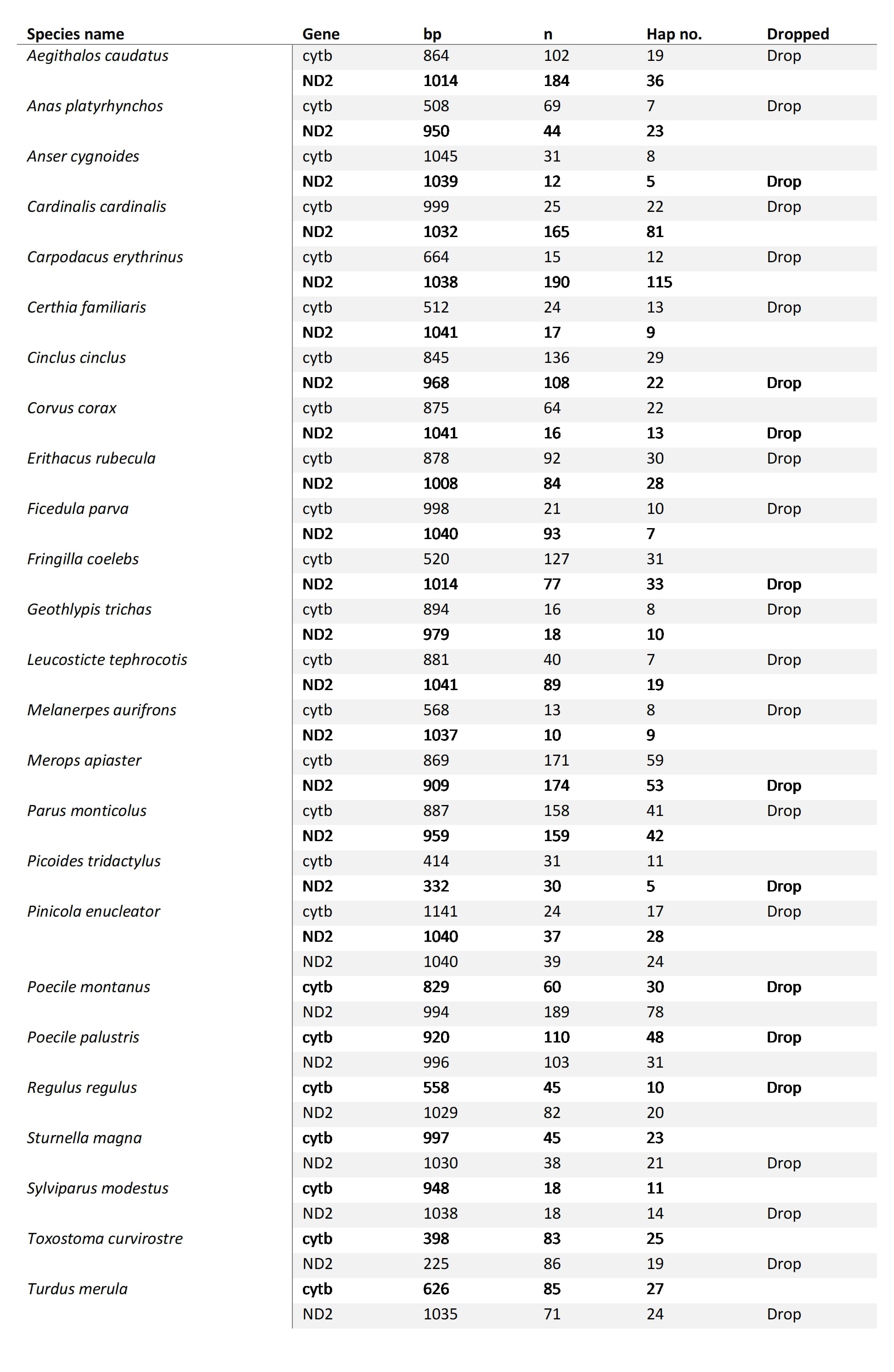

Supplementary Table 1. 25 species included in analysis for which sequence data was available for both cytb and ND2.
