## Supplementary material for "mtDNA-based reconstructions of change in effective population sizes of Holarctic birds do not agree with their reconstructed range sizes based on paleoclimates": Pipeline for SDM analyses

### SDM analyses of Holarctic birds

*Michela Leonardi, Department of Zoology, University of Cambridge.*

*2019-08-07*

#### Introduction

The present document contains the R code used for performing SDM analyses (section *Analyses*) and plotting the outputs (section *Plots*) for Miller et al. "mtDNA-based reconstructions of change in effective population sizes of Holarctic birds do not agree with their reconstructed range sizes based on paleoclimates". The two sections of the document can be run as separate scripts.

#### Analyses

##### Prepare the environment

Setting the working directory and variables, load a low resolution world map.

```
setwd("C:/Users/miche/Desktop/SDM Birds/Holarctic birds 2019_08_07")

# libraries
library(rgbif)
library(rworldmap)
library(data.table)
library(PBSmapping)
library(rgeos)
library(maptools)
library(beanplot)
library(raster)
library(rgdal)
library(SDMTools)
library(RColorBrewer)
library(caret)
library(biomod2)
library(spThin)
library(blockCV)

# Function to modify coordinates
wherenearest <- function(val,matrix) {
  dist=abs(matrix-val)
  index=which.min(dist)
  return( index )
}

# Load world map
WorldMap <- getMap(resolution="low")

# vector of variable names
vars <- c("npp","lai","BIO1","BIO4","BIO5","BIO6","BIO7","BIO8","BIO9","BIO10",
          "BIO11","BIO12","BIO13","BIO14","BIO15","BIO16","BIO17","BIO18","BIO19")
```

```
# time for past projections (thousands of years ago)
kyr <- 21
```

#### Analyses

In order to succesfully perform the analyses please please move the "Species.csv" file from the "Input_20190807" folder into the working directory. The code below performs on each species the following tasks (each one marked by a numbered title within the code):

1. Loading data
2. Data preparation
3. Violin plots
4. Cross-correlation between variables
5. Thinning
6. Preparing the environment for the biomod2 analyses
7. Pseudo-absence selection
8. Formatting the species input file for biomod2
9. Create the biomod2 input object
10. Spatial block cross-validation
11. Model calibration
12. Projection to the entire study area
13. Ensemble modelling
14. Validation of the model
15. Ensemble forecasting
16. Projection backwards in time

```
# read species data
csv <- read.table("Species.csv",sep=",", header=TRUE)

for (i in 1:dim(csv)[1]){

  #==================#
  # 1. Loading data #
  #==================#

  # variables
  sp <- as.character(csv$Species[i])
  filename <- as.character(csv$GBIF.code[i])
  biome <- as.character(csv$biome[i])
  region <- as.character(csv$region[i])
  outpath <- paste(sp, "/", region,"/", sep="")
  dir.create(file.path(outpath),recursive=TRUE)

  # Species name with "_" or "." instead of space
  sp_name <- paste(unlist(strsplit(sp, " "))[1],
                   unlist(strsplit(sp, " "))[2], sep="_")
  sp.name <- paste(unlist(strsplit(sp, " "))[1],
                   unlist(strsplit(sp, " "))[2], sep=".")

  # define limit coordinates (W,E,S,N) and plot width
  if (region=="NAmerica"){
    coord <- c(-180,-15,8,90)
    wtd <- 10
    h <- 8
```

```r
} else if (region=="Eurasia"){
  coord <- c(-12,170,5,85)
  wtd <- 13
  h <- 6
} else {
  stop('Region name is neiter "Eurasia" or "NAmerica", please check spelling')
}

# define colors (filled and semi-transparent)
if (biome=="forest"){
  color <- rgb(0.55,0.75,0.45,1)
  Tcolor <- rgb(0.55,0.75,0.45,0.75)
} else if (biome=="grassland"){
  color <- rgb(0,0.83,0,1)
  Tcolor <- rgb(0,0.83,0,0.6)
} else if (biome=="shrubland"){
  color <- rgb(0.65,0.5,0.2,1)
  Tcolor <- rgb(0.65,0.5,0.2,0.45)
} else if (biome=="wetland"){
  color <- rgb(0.4,0.7,0.8,1)
  Tcolor <- rgb(0.4,0.7,0.8,0.75)
} else if (biome=="other"){
  color <- rgb(0.5,0.7,0.7,1)
  Tcolor <- rgb(0.5,0.7,0.7,0.75)
} else {
  stop('Biome name is neither "forest", "grassland", "shrubland", "wetland", or "other"
       please check spelling')
}

# Environmental variables for the whole area, in the present
envdata <- read.table(paste("Input_20190807/", region, "/EnvirVar/EnvirVar_",
                            region,"_0.txt", sep=""),
                      header=TRUE, sep=" ")
colnames(envdata)[1:2] <- c("long","lat")

#Extract latitude and longitude
lon <- as.vector(unique(envdata$long))
lat <- as.vector(unique(envdata$lat))

#======================#
# 2. Data preparation #
#======================#

# download GBIF file
occ_download_get(filename, overwrite=TRUE)

# unzip file
unzip(paste(filename,".zip",sep=""), overwrite=TRUE,
      exdir=".", unzip="internal")

# read file
distrib <- fread(paste(filename,".csv",sep="")) #GBIF file
```

```
# filter by latitude and longitude
distrib <- distrib[distrib$decimalLongitude > min(lon) &
            distrib$decimalLongitude < max(lon) &
            distrib$decimalLatitude > min(lat) &
            distrib$decimalLatitude < max(lat),]

# filter by coordinate uncertainty, and only keep relevant columns
distrib <- distrib[distrib$coordinateUncertaintyInMeters < 10000 |
             is.na(distrib$coordinateUncertaintyInMeter),
           c("gbifID","decimalLatitude","decimalLongitude")]

# modify latitude and longitude to match the grid of the climatic reconstructions
distrib$decimalLatitude <- sapply(distrib$decimalLatitude, function(x)
  lat[wherenearest(x,lat)])
distrib$decimalLongitude <- sapply(as.numeric(distrib$decimalLongitude), function(x)
  lon[wherenearest(x,lon)])

# remove duplicates
distrib <- distrib[!duplicated(distrib[,c("decimalLatitude","decimalLongitude")]),]

# read mask data
SpeciesMask<-readShapePoly(paste("Input_20190807/Masks/", sp_name, sep=""),
                         proj4string=CRS("+proj=longlat +datum=WGS84"))

sapply(slot(SpeciesMask, "polygons"), function(x) slot(x, "ID"))

# subset it for presence=1 or 2
SpeciesMask.sub<-SpeciesMask[SpeciesMask$PRESENCE<3,]
SpeciesMask.sub<-SpeciesMask[SpeciesMask$ORIGIN<3,]

# select the summer/resident component
SpeciesMask<-SpeciesMask.sub[SpeciesMask.sub$SEASONAL%in%c("1","2"),]

# create Polysets for summer and resident component:
# merging all polygons to create a single PID, needed for later operations
SpeciesMask<-SpatialPolygons2PolySet(SpeciesMask)
if (length(unique(SpeciesMask$PID))>1) {
  SpeciesMask2<-joinPolys(SpeciesMask,operation="UNION")
}
SpeciesMask<-PolySet2SpatialPolygons(SpeciesMask)

# Plot map with observations and mask
# plot(WorldMap, col="cornsilk", bg="lightblue1", border="grey",
# xlim=coord[1:2], ylim=coord[3:4], lwd=0.05)
# points(as.numeric(distrib$decimalLongitude),as.numeric(distrib$decimalLatitude),
# pch="*", col=color)
# plot(SpeciesMask, add=TRUE)

# Subset data if within polygon
# define coordinates
xy <- distrib[,c(3,2)]

#transform into SpatialPointsDataFrame
```

```r
df <- SpatialPointsDataFrame(coords=xy, data=distrib[,1],
                             proj4string=SpeciesMask@proj4string)

# keep only points in shapefile
distrib <- df[!is.na(over(df,SpeciesMask)),]

# create file with observations (3 columns: long, lat, observation - 1 as presence)
distrib <- cbind(distrib$decimalLongitude, distrib$decimalLatitude,
                 rep(1, length(distrib$decimalLongitude)))

colnames(distrib) <- c("long","lat",sp_name)

# save file
write.table(distrib, paste(outpath,sp_name,"_distrib.txt", sep=""))

#=================#
# 3. Violin plots #
#=================#

# Remove NA from environmental data
envdata <-  envdata[complete.cases(envdata), ]

# Extract variables for observations
obs <- merge(distrib[,c(1,2)], envdata)
#head(obs)

# merge data in a table, last column distinguish between baseline and observations
envdata[,"set"] <- "baseline"
obs[,"set"] <- "obs"
data <- rbind(envdata,obs)

# Variable distribution plot
png(paste(outpath, sp_name,"_variables.png", sep=""),
    height=9, width=8,  units='in', res=600)
par(mfrow=c(4, 5),
    oma=c(0, 1, 5, 0),
    mar=c(1, 1, 3, 1),
    mgp=c(0.5, 0.5, 0))

# for each environmental variable
for (v in vars){

  # Beanplot distributions
  beanplot(data[data$set=="baseline",v],data[data$set=="obs",v],
           bw="nrd",side="both", col=list("black",color),
           border=c("black", color), what=c(1,1,1,0),
           main=v ,xaxt='n')
}
plot.new()
legend("center", c("Baseline", "Species\noccurrences"), fill=c("black", color),
       border=NA, bty="n", cex=1.5)

title(main=paste(sp,"climatic variables", sep=" "),
```

```r
      cex.main= 3, outer=TRUE, line=2)
dev.off()

#================================#
# 4. Variables cross-correlation #
#================================#

# function to plot, courtesy of Raquel A. Garcia, Stellenbosch University, South Africa.
panel.cor <- function(x, y, digits=2, prefix="", cex.cor)
{
  usr <- par("usr"); on.exit(par(usr))
  par(usr=c(0, 1, 0, 1))
  r <- abs(cor(x, y))
  txt <- format(c(r, 0.123456789), digits=digits)[1]
  txt <- paste(prefix, txt, sep="")
  if(missing(cex.cor)) cex <- 0.8/strwidth(txt)

  test <- cor.test(x,y)
  # borrowed from printCoefmat
  Signif <- symnum(test$p.value, corr=FALSE, na=FALSE,
                   cutpoints=c(0, 0.001, 0.01, 0.05, 0.1, 1),
                   symbols=c("***", "**", "*", ".", " "))

  text(0.5, 0.5, txt, cex=cex * r)
  text(.8, .8, Signif, cex=cex, col=2)
}

# Pairwise correlation matrix
cormat <- cor(envdata[,vars])

# plot correlation for all variables
png(filename=paste(outpath, sp_name, "_correlation_all_vars.png",sep=""),
    width=1200, height=900)
pairs(envdata[,vars], lower.panel=panel.smooth, upper.panel=panel.cor)
dev.off()

# define uncorrelated variables
uncor <- findCorrelation(cormat, cutoff=0.7)

# plot correlation for chosen variables
png(filename=paste(outpath, sp_name,"_correlation_uncorr_vars.png", sep=""),
    width=1200, height=900)
pairs(envdata[,vars[-c(uncor)]], lower.panel=panel.smooth, upper.panel=panel.cor)
dev.off()

#=============#
# 5. Thinning #
#=============#

# thinning
t <- thin(loc.data=as.data.frame(distrib), lat.col="lat", long.col="long",
          spec.col=sp_name, thin.par=70, reps=100, locs.thinned.list.return=TRUE,
          write.files=TRUE, max.files=10, out.dir=paste(outpath,"Thin/", sep=""),
```

```r
          out.base=sp_name, write.log.file=FALSE)

ThinDistrib <- read.table(paste(outpath, "Thin/", sp_name, "_thin1.csv", sep=""),
                          header=TRUE,  sep=",")
ThinDistrib <- ThinDistrib[,c(2,3,1)]

# plot thinning
png(paste(outpath, sp_name, "_thinned_dataset.png", sep=""),
    height=6, width=wtd,  units='in', res=600)
par(mfrow=c(1, 2),
    oma=c(0, 0, 5, 0),
    mar=c(3, 1, 3, 1),
    mgp=c(2, 1, 0))

plot(WorldMap, col="cornsilk", bg="lightblue1", border="grey", xlim=coord[1:2],
     ylim=coord[3:4], lwd=0.05, main=paste("Dataset\nN =", dim(distrib)[1], sep=" "))
points(as.numeric(distrib[,"long"]),as.numeric(distrib[,"lat"]), pch="*", col=color)

plot(WorldMap, col="cornsilk", bg="lightblue1", border="grey",
     xlim=coord[1:2], ylim=coord[3:4], lwd=0.05,
     main=paste("Thinned dataset\nN =", dim(ThinDistrib)[1], sep=" "))
plot(SpeciesMask, add=TRUE, col=color, border=NA)
points(as.numeric(ThinDistrib$long), as.numeric(ThinDistrib$lat), pch="*", col="black")

title(main=paste(sp,"occurrences", sep=" "),cex.main= 3,
      outer=TRUE, line=2)

dev.off()

#===========================================#
# 6. Preparing the environment for biomod2 #
#===========================================#

# Raster maps for climatic variables through time
basefile <- paste("Input_20190807/", region, "/", sep="")

# Color palette
c1 <- colorRampPalette(c("khaki1",color))(6)
c2 <- colorRampPalette(c(color,"black"))(8)
cols <- c(c1[2:5],c2[2:8])

# environmental variables
expl.var <- stack()

vars1 <- vars[-c(uncor)]
# create raster stack
for(v in 1:length(vars1)){
  r <- raster(paste(basefile, vars1[v], "/", region, "_",vars1[v],"_0" ,".grd", sep=""),
              RAT=FALSE)
  expl.var <- stack( expl.var, r)
}

#=============================#
```

```r
# 7. Pseudo-absence selection #
#==============================#

# background data
bg <- data.frame(cbind(envdata[,1:2], species=rep(0,dim(envdata)[1])))
colnames(bg)[3] <- sp.name

# define coordinates
xy <- bg[,1:2]

#transform into SpatialPointsDataFrame
df <- SpatialPointsDataFrame(coords=xy, data=bg,
                             proj4string=SpeciesMask@proj4string)

# remove points within the mask
bg.clean <- df[is.na(over(df,SpeciesMask)),]
bg.clean <- data.frame(bg.clean[,1:3])

# number of absences and presences
pres <- dim(ThinDistrib)[1]
abs <- dim(ThinDistrib)[1]

# Absences table
abs.table <- data.frame(matrix("FALSE",abs*5,7), stringsAsFactors=FALSE)
colnames(abs.table) <- c("long","lat",paste("RUN",c(1:5),sep=""))

# variable to define the first line to be written
start <- 1

for (k in 1:5){
  index <- sample(1:dim(bg.clean)[1], abs)
  abs.table[seq(start,abs*k),1:2] <- bg.clean[index,1:2]
  abs.table[seq(start,abs*k),2+k] <- rep("TRUE",abs)
  start <- (abs*k)+1
}

#===================================#
# 8. Formatting species input file #
#===================================#

# species presences + background
resp.var <- as.numeric(c(rep(1, pres),rep(0,abs*5)))
# coordinates
resp.xy <- rbind(ThinDistrib[,c("long","lat")], abs.table[,c("long","lat")])

# add presences to PA.table
pres.table <- data.frame(cbind(ThinDistrib[,c("long","lat")]), matrix("TRUE",pres,5))
colnames(pres.table) <- c("long","lat",paste("RUN",c(1:5),sep=""))

# merge tables for presences and pseudoabsences
PA.table <- rbind(pres.table[,-c(1,2)], abs.table[,-c(1,2)])
PA.table[] <- lapply(PA.table, as.logical)
```

```
#===================================#
# 9. Creating biomod2 input object #
#===================================#

biomodData <- BIOMOD_FormatingData(resp.var=resp.var,     # species distribution
                                   expl.var=expl.var,     # environmental variables
                                   resp.xy=resp.xy,       # coordinates of species
                                   resp.name=sp.name,     # name of the species column
                                   PA.strategy="user.defined",# pseudo absences
                                   PA.table=PA.table,
                                   na.rm=TRUE)             # do not consider NA

png(paste(outpath, sp_name, "_pseudoabsences_datasets.png", sep=""),
    width=7, height=7, units='in', res=600)
plot(biomodData)
dev.off()

# To store the resulting object
save(biomodData, file=paste(outpath, sp_name, "_BiomodData", sep=""))

#=====================================#
# 10. Spatial block cross-validation #
#=====================================#

# presences and absences taken from biomodData (with NA removed)
resp <- cbind(biomodData@coord,sp=biomodData@data.species)
resp[] <- lapply(resp, as.numeric)
colnames(resp)[3] <- sp

# transform into SpatialPointsDataFrame
resp <- SpatialPointsDataFrame(resp[,c("long", "lat")], resp, proj4string=crs(expl.var))

# create spatial blocks and saves plot
png(paste(outpath,sp_name,"_spatial_blocks.png",sep=""),
    height=6, width=10,  units='in', res=600)

# If NAmerica horizontal blocks, if Eurasia vertical blocks
if (region=="NAmerica"){
  sb <- spatialBlock(speciesData=resp,
                     species=sp,
                     rasterLayer=expl.var,
                     rows=15,
                     k=5,
                     selection="systematic",
                     biomod2Format=TRUE)

} else if (region=="Eurasia") {
  sb <- spatialBlock(speciesData=resp,
                     species=sp,
                     rasterLayer=expl.var,
                     cols=15,
                     k=5,
                     selection="systematic",
```

```
                     biomod2Format=TRUE)
}
dev.off()

# !!!!!!!!!!!!!!!!!!!!!!!!!!!!!!!!!!!!!!!!!!!!!!!!!!!!!!!!!!!!! #
# WARNING:                                                        #
# it is necessary to check if all the runs contain presences #
# (>10) and absences. If not the table must be subset within #
# the BIOMOD_Modeling function (see next warning, line 491)  #
# !!!!!!!!!!!!!!!!!!!!!!!!!!!!!!!!!!!!!!!!!!!!!!!!!!!!!!!!!!!!! #

DataSplitTable <- sb$biomodTable

# add coordinates to the table
dst.xy <- cbind(resp$long,resp$lat,DataSplitTable)
colnames(dst.xy)[1:2] <- c("x", "y")

# plot
png(paste(outpath, sp_name,"_blocks_CV.png", sep=""), width=7, height=h,
    units='in', res=600)
par(mfrow=c(3, 2),
    oma=c(0, 0, 5, 0),
    mar=c(1, 1, 1, 1),
    mgp=c(2, 1, 0))

for (k in c(1:5)){

  col <- DataSplitTable[,paste("RUN",k,sep="")]
  col[col=="FALSE"] <- color
  col[col=="TRUE"] <- "black"

  plot(WorldMap, col="cornsilk", bg="lightblue1", lwd=0.05, border="grey",
       xlim=c(min(lon), max(lon)), ylim=c(min(lat), max(lat)),
       main=paste("RUN",k,sep=""))
  points(as.numeric(dst.xy[,"x"]), as.numeric(dst.xy[,"y"]), pch="*", col=col)
}

# title
title(main=paste(sp," spatial blocks cross-validation",sep=""), cex.main=2, outer=TRUE,
      line=3)

dev.off()

#=======================#
# 11. Model calibration #
#=======================#

# Model parameters
ModOptions <- BIOMOD_ModelingOptions()

mods <- c("GLM","GBM","GAM","RF") # GBM=boosted regression tree

ModelOut <- BIOMOD_Modeling(biomodData,               # input data
```

```
                          models=mods,                  # algorithms
                          models.options=ModOptions,    # options
# !!!!!!!!!!!!!!!!!!!!!!!!!!!!!!!!!!!!!!!!!!!!!!!!!!!!!!!!!!! #
# WARNING:                                                      #
# in DataSplitTable only consider columns (=splits)          #
# including BOTH presences (>10) and absences                #
# !!!!!!!!!!!!!!!!!!!!!!!!!!!!!!!!!!!!!!!!!!!!!!!!!!!!!!!!!!! #
                          DataSplitTable=DataSplitTable,# DataSplitTable issued by blockCV
                          models.eval.meth=c("TSS"),    # method for evaluation
                          SaveObj=T,                    # save object
                          modeling.id=paste(sp.name,"_modelOut", sep=""),
                          do.full.models=FALSE,
                          rescal.all.models=T)

# model evaluations
ModelEvaluation <- get_evaluations(ModelOut)
ModelEvaluation["TSS","Testing.data",,,]

# TSS scores
write.table(ModelEvaluation["TSS","Testing.data",,,],
            paste(outpath, sp_name, "_models_eval.txt", sep=""))

#==========================================#
# 12. Projecting to the entire study area #
#==========================================#

new.env <- expl.var
Projection <- BIOMOD_Projection(modeling.output=ModelOut,    # model
                             selected.models="all",     # models to select
                             new.env= new.env,          # environmental data to project to
                             proj.name=region,
                             binary.meth=c("TSS"),      # method for binary transformation
                             filtered.meth=c("TSS"),    # method to set values below zero
                             build.clamping.mask=T,     # out-of-calibration range
                             compress="xz")

# # make some plots sub-selected by str.grep argument
# png(paste(outpath,sp_name,"_proj_RF_RUN1.png", sep=""),
#     width=9, height=12, units='in', res=600)
# plot(Projection, str.grep='RF')
# dev.off()

#=========================#
# 13. Ensemble modelling #
#=========================#

Ensemble <- BIOMOD_EnsembleModeling(modeling.output=ModelOut,
                                    chosen.models="all",       # models used
                                    em.by="all",               # by algorithm
                                    eval.metric=c("TSS"),      # evaluation metric
                                    eval.metric.quality.threshold=c(0.7),
                                    models.eval.meth=c("TSS"), # evaluation method
                                    prob.mean=T,
```

```
                                    prob.median=T,
                                    committee.averaging=T,
                                    prob.mean.weight=T,
                                    prob.mean.weight.decay="proportional")

#==============================#
# 14. Validation of the model #
#==============================#

# evaluation scores for the ensemble model
EnsembleEvaluation <- get_evaluations(Ensemble)
write.table(EnsembleEvaluation, file=paste(outpath, sp_name,"_Ensemble_eval.txt", sep=""))

# change dimension names to make them more easily accessible
names(EnsembleEvaluation) <- c("mean_TSS","median_TSS","ca_TSS","wmean_TSS")

# Create a table with evaluation scores, sensitivity and specificity
out<- rbind(c(EnsembleEvaluation$mean_TSS[,1], EnsembleEvaluation$median_TSS[,1],
              EnsembleEvaluation$ca_TSS[,1], EnsembleEvaluation$wmean_TSS[,1]),
            c(EnsembleEvaluation$mean_TSS[,3], EnsembleEvaluation$median_TSS[,3],
              EnsembleEvaluation$ca_TSS[,3], EnsembleEvaluation$wmean_TSS[,3])/100,
            c(EnsembleEvaluation$mean_TSS[,4], EnsembleEvaluation$median_TSS[,4],
              EnsembleEvaluation$ca_TSS[,4], EnsembleEvaluation$wmean_TSS[,4])/100)

# index for the maximum TSS
maxTSS <- which.max(out[1,])

# Plot barplots of evaluation
colnames(out) <- c("Mean","Median","Committee\naverage","Weighted\nmean")
rownames(out) <- c("Evaluation score", "Sensitivity","Specificity")

png(paste(outpath, sp_name,"_evaluation_scores_ensemble.png", sep=""),
    width=6, height=4.5, units='in', res=600)
layout(matrix(c(1,2), ncol=1, byrow=TRUE), heights=c(3.8,0.7))
par(oma=c(0, 0, 4, 0), # rows of text at the outer bottom left top right margin
    mar=c(2, 3, 3, 1))#, # space for row of text at ticks and to separate plots

barplot(t(out), beside=TRUE, col=cols[c(6,2,8,4)], border=NA, ylim=c(0,1))

par(mai=c(0,0,0,0))
plot.new()
legend("center", colnames(out), fill=cols[c(6,2,8,4)], border=NA, bty="n",ncol=4)

title(main=paste("Evaluation scores -", sp, sep=" "), cex.main=1.5, outer=TRUE, line=1)

dev.off()

#==========================#
# 15. Ensemble forecasting #
#==========================#

EnsembleProjection <- BIOMOD_EnsembleForecasting(EM.output=Ensemble,# output from ensemble
                                projection.output=Projection,  # output from projection
```

```
                                selected.models="all",          # model chosen
                                binary.meth=c("TSS"),           # method for binary transf
                                filtered.meth=c("TSS"),         # method for filtering
                                compress=T)                     # compression

#list of files in directory
filelist <- list.files(paste(sp.name,"/proj_",region,"/", sep=""))
# selecting consensus projections present
enslist <- filelist[grep("ensemble", filelist)]
PresEnsemble <- stack(paste(sp.name,"/proj_",region,"/", enslist[2], sep=""))

EnsembleMethods <- c("Mean", "Median", "Committee average", "Weighted mean")

# plot without observation points
png(paste(outpath, sp_name,"_ensemble_projection.png", sep=""),
    width=12, height=10, units='in', res=600)
par(mfrow=c(2, 2),
    oma=c(0, 0, 4, 0),
    mar=c(3, 3, 2, 5),
    mgp=c(2, 1, 0))

for (x in 1:4) {
  #plot selected rasters (ensemble models)
  plot(raster(PresEnsemble,x),
       main=EnsembleMethods[x])
}
title(main=paste("Ensemble projections -", sp, sep=" "), cex.main= 1.5,
      outer=TRUE, line=1)

dev.off()

# plot with observation points
png(paste(outpath, sp_name,"_ensemble_projection_points.png", sep=""),
    width=12, height=10, units='in', res=600)
par(mfrow=c(2, 2),
    oma=c(0, 0, 4, 0),
    mar=c(3, 3, 2, 5),
    mgp=c(2, 1, 0))

for (x in 1:4) {
  #plot selected rasters (ensemble models)
  plot(raster(PresEnsemble,x),
       main=EnsembleMethods[x])
  points(ThinDistrib[,"long"], ThinDistrib[,"lat"], pch="*")
}

title(main=paste("Ensemble projections -", sp, sep=" "), cex.main= 1.5,
      outer=TRUE, line=1)

dev.off()

#==================================#
# 16. Projection backwards in time #
```

```r
#==================================#

# environmental variables
past.env <- stack()

# create raster stack
for(v in 1:length(vars1)){
  r <- raster(paste(basefile, vars1[v], "/", region, "_",vars1[v],"_", kyr ,".grd", sep=""),
              RAT=FALSE)
  past.env <- stack(past.env, r)
}

# Project model into the past
PastProjection <- BIOMOD_Projection(modeling.output=ModelOut,   # model
                          selected.models="all",      # model to select
                          new.env= past.env,          # environmental data to project to
                          proj.name=paste(kyr,region, sep="_"),
                          binary.meth="TSS",          # method for binary transformation
                          filtered.meth="TSS",        # method to set values below zero
                          build.clamping.mask=T,      # out-of-calibration range
                          compress="xz")

# Project ensemble to the past
PastEnsemble <- BIOMOD_EnsembleForecasting(EM.output=Ensemble,     # output from ensemble
                              projection.output=PastProjection,  # output from projection
                              selected.models="all",       # model chosen
                              binary.meth=c("TSS"),        # eval method for bin transf
                              filtered.meth=c("TSS"),      # eval method for filtering
                              compress=T)                  # compression?


filelistP <- list.files(paste(sp.name,"/proj_",kyr,"_", region, "/", sep=""))
enslist <- filelistP[grep("ensemble", filelistP)]  # selecting consensus projections
PastEnsemble <- stack(paste(sp.name, "/proj_",kyr,"_", region, "/", enslist[2], sep=""))

# plot all rasters
png(paste(outpath, sp_name,"_",kyr,"_all_ensemble_projections.png", sep=""),
    width=12, height=9, units='in', res=600)

op<-par(mfrow=c(2, 2),
        oma=c(0, 0, 4, 1),
        mar=c(3, 3, 2, 5),
        mgp=c(2, 1, 0))

for (x in 1:4) {
  #plot selected rasters (ensemble models)
  plot(raster(PastEnsemble,x),
       main=EnsembleMethods[x])
}
title(main=paste("Ensemble projections LGM -", sp, sep=" "), cex.main= 1.5,
      outer=TRUE, line=1)

dev.off()
```

```
  #create LGM directory
  dir.create(paste(outpath,"LGM",sep=""), showWarnings=FALSE)

  # one plot for each statistic
  for (x in 1:4) {
    png(paste(outpath, "LGM/", sp_name, "_", kyr, "_", EnsembleMethods[x],
              "_ensemble_projection.png", sep=""),
        width=12, height=9, units='in', res=600)
    plot(raster(PastEnsemble,x),
         main=paste(sp, kyr, "kya", EnsembleMethods[x], "ensemble projection", sep=" "))
    dev.off()
  }

  # index of the layers to use

  for (j in 1:4){
    # present
    PresRaster <- raster(PresEnsemble,j)
    writeRaster(PresRaster, paste(outpath, "LGM/", sp_name, "_", EnsembleMethods[j],
                                  "_pres", sep=""),
                overwrite=TRUE)

    # past
    PastRaster <- raster(PastEnsemble,j)
    writeRaster(PastRaster, paste(outpath, "LGM/", sp_name, "_", EnsembleMethods[j],
                                  "_LGM", sep=""),
                overwrite=TRUE)

    # difference between the two (present - past)
    difference <- PresRaster-PastRaster
    writeRaster(difference, paste(outpath, "LGM/", sp_name, "_", EnsembleMethods[j],
                                  "_pres-past", sep=""),
                overwrite=TRUE)
  }
  # write on file
  # clean dataset
  csv[i,"Clean.dataset"] <- dim(distrib)[1]
  # thinned dataset
  csv[i,"Thinned.dataset"] <- dim(ThinDistrib)[1]

  # Overwrite species file including results
  write.csv(csv, file="Species.csv", quote=FALSE, row.names=FALSE)

}
```

### Plots

#### Prepare environment

Please consider that with respect to the SDM analysis script some files and folders have been moved: the output folder from biomod called *"Genus.species"* has been moved in the *"Genus species"* folder within "Holarctic birds 2019_08_07/Results_20190807" (set as working directory) that must also contain the "Species.csv" file issued by the previous script.

```r
setwd("C:/Users/miche/Desktop/SDM Birds/Holarctic birds 2019_08_07/Results_20190807")


# libraries
library(rgbif)
library(rworldmap)
library(data.table)
library(PBSmapping)
library(rgeos)
library(maptools)
library(beanplot)
library(raster)
library(rgdal)
library(SDMTools)
library(RColorBrewer)
library(caret)
library(biomod2)
library(spThin)
library(blockCV)

# Function to modify coordinates
wherenearest <- function(val,matrix) {
  dist=abs(matrix-val)
  index=which.min(dist)
  return( index )
}

# Load world map
WorldMap <- getMap(resolution="low")

# time for past projections (thousands of years ago)
kyr <- 21

# read species data
csv <- read.table("Species.csv",sep=",", header=TRUE)

# Colors for overlap map
cols <- c( "cornsilk",
           brewer.pal(9,"YlGn")[4],
           brewer.pal(9,"Blues")[6],
           "darkslategray"
)
}
```

#### Plots

The code below performs on each species the following tasks (each one marked by a numbered title within the code).

1. Loading data
2. Calculate the areas
3. Calculate the overlap between present and LGM
4. Create a multiplot including observations, thinning, present day, LGM
5. Plot of the overlap between the present and the LGM

```r
for (i in c(1:dim(csv)[1])){

  #==================#
  # 1. Loading data #
  #==================#

  # variables
  sp <- as.character(csv$Species[i])
  biome <- as.character(csv$biome[i])
  region <- as.character(csv$region[i])
  outpath <- paste(sp, "/", region,"/", sep="")

  # Species name with "_" or "." instead of space
  sp_name <- paste(unlist(strsplit(sp, " "))[1],
                   unlist(strsplit(sp, " "))[2], sep="_")
  sp.name <- paste(unlist(strsplit(sp, " "))[1],
                   unlist(strsplit(sp, " "))[2], sep=".")

  # define limit coordinates (W,E,S,N) and plot width
  if (region=="NAmerica"){
    coord <- c(-180,-15,8,90)
    wtd <- 9.5
    h <- 8
  } else if (region=="Eurasia"){
    coord <- c(-12,170,5,85)
    wtd <- 13
    h <- 6
  } else {
    stop('Region name is neiter "Eurasia" or "NAmerica", please check spelling')
  }

  # define colors (filled and semi-transparent)
  if (biome=="forest"){
    color <- rgb(0.55,0.75,0.45,1)
    Tcolor <- rgb(0.55,0.75,0.45,0.75)
  } else if (biome=="grassland"){
    color <- rgb(0,0.71,0,1)
    Tcolor <- rgb(0,0.71,0,0.6)
  } else if (biome=="shrubland"){
    color <- rgb(0.65,0.5,0.2,1)
    Tcolor <- rgb(0.65,0.5,0.2,0.45)
  } else if (biome=="wetland"){
    color <- rgb(0.4,0.7,0.8,1)
    Tcolor <- rgb(0.4,0.7,0.8,0.75)
  } else if (biome=="other"){
    color <- "slategray"
    Tcolor <-"slategray"
  } else {
    stop('Biome name is neither "forest", "grassland", "shrubland", "wetland", or "other".
         please check spelling')
  }

  # Environmental variables for the whole area, in the present
```

```r
envdata <- read.table(
  paste("C:/Users/miche/Desktop/SDM Birds/Holarctic birds 2019_08_07/Input_20190807/",
        region, "/EnvirVar/EnvirVar_", region,"_0.txt", sep=""), header=TRUE, sep=" ")
colnames(envdata)[1:2] <- c("long","lat")

#Extract latitude and longitude
lon <- as.vector(unique(envdata$long))
lat <- as.vector(unique(envdata$lat))

# read mask data
SpeciesMask<-readShapePoly(
  paste("C:/Users/miche/Desktop/SDM Birds/Holarctic birds 2019_08_07/Input_20190807/Masks/",
        sp_name, sep=""), proj4string=CRS("+proj=longlat +datum=WGS84"))

sapply(slot(SpeciesMask, "polygons"), function(x) slot(x, "ID"))

# subset it for presence=1 or 2
SpeciesMask.sub<-SpeciesMask[SpeciesMask$PRESENCE<3,]
SpeciesMask.sub<-SpeciesMask[SpeciesMask$ORIGIN<3,]

# select the summer/resident component
SpeciesMask<-SpeciesMask.sub[SpeciesMask.sub$SEASONAL%in%c("1","2"),]

# create Polysets for summer and resident component:
# merging all polygons to create a single PID, needed for later operations
SpeciesMask<-SpatialPolygons2PolySet(SpeciesMask)
if (length(unique(SpeciesMask$PID))>1) {
  SpeciesMask2<-joinPolys(SpeciesMask,operation="UNION")
}
SpeciesMask<-PolySet2SpatialPolygons(SpeciesMask)

# read file with cleaned observations
distrib <- read.table(paste(outpath,sp_name,"_distrib.txt", sep=""), header=TRUE)

# read file with thinned observations
ThinDistrib <- read.table(paste(outpath, "Thin/", sp_name, "_thin1.csv", sep=""),
                          header=TRUE, sep=",")
ThinDistrib <- ThinDistrib[,c(2,3,1)]

# read file with model evaluation and get maximum
EnsembleEvaluation <- read.table(paste(outpath, sp_name, "_Ensemble_eval.txt", sep=""),
                                 header=TRUE)
tss <- EnsembleEvaluation[1,c(1,5,9,13)]
maxTSS <- which(tss==max(tss))[1]
EnsembleMethods <- c("Mean", "Median", "Committee average", "Weighted mean")

# read file for the present and create rasters for stats of interest
# list of files in the directory
filelist <- list.files(paste(sp,"/",sp.name,"/proj_",region,"/", sep=""))
# selecting consensus projections
enslist <- filelist[grep("ensemble", filelist)]
PresEnsemble <- stack(paste(sp,"/",sp.name,"/proj_", region, "/", enslist[2], sep=""))
```

```r
# read file for the past
filelist <- list.files(paste(sp,"/",sp.name,"/proj_",kyr,"_",region,"/", sep=""))
enslist <- filelist[grep("ensemble", filelist)]
PastEnsemble <- stack(paste(sp,"/",sp.name,"/proj_",kyr,"_",region,"/",
                            enslist[2], sep=""))

#==========#
# 2. Areas #
#==========#

# binary transformation raster present
FlatPresRaster <- BinaryTransformation(raster(PresEnsemble,maxTSS),500)
# project to Eckert IV equal-area pseudocylindrical projection (grid size=100km)
PresRaster <- FlatPresRaster
crs(PresRaster) <- CRS('+init=EPSG:4326')
PresRaster <- projectRaster(PresRaster,crs="+proj=eck4",over=T,method="ngb", res=50000)
# Calculate number of cells
PresArea <- tapply(PresRaster, PresRaster[], sum)["1"]
# Area in square km
PresAreaKm <- PresArea*2500

# binary transformation raster past
FlatPastRaster <- BinaryTransformation(raster(PastEnsemble,maxTSS),500)
# project to Eckert IV equal-area pseudocylindrical projection
PastRaster <- FlatPastRaster
crs(PastRaster) <- CRS('+init=EPSG:4326')
PastRaster <- projectRaster(PastRaster,crs="+proj=eck4",over=T,method="ngb",res=50000)
# Calculate number of cells
PastArea <- tapply(PastRaster, PastRaster[], sum)["1"]
# Area in square km
PastAreaKm <- PastArea*2500

# write on file
# area LGM
csv[i,"LGM"] <- PastAreaKm
# area present
csv[i,"Present"] <- PresAreaKm

#========================#
# 3. Overlap present-LGM #
#========================#

# sum
PastRaster[is.na(PastRaster)] <- 0
OverlapRaster <- PresRaster+(PastRaster*2)
# Calculate number of cells
PresPastAreas <- tapply(OverlapRaster, OverlapRaster[], sum)/(c(1,1,2,3))

# overlap index (how much of the present range is suitable during LGM)
overlap <- (sum(PresPastAreas[c("3")])/sum(PresPastAreas[c("1","3")]))

# write on file
csv[i, "Pres.LGM.Overlap"] <- overlap
```

```r
#====================================================#
# 4. Multiplot observations, thinning, present, LGM #
#====================================================#

# Ice
PresIce <- raster(
  'C:/Users/miche/Desktop/SDM Birds/Holarctic birds 2019_08_07/Input_20190807/Mask_0.nc',
                  var="ice_thickness")
PastIce <- raster(
  'C:/Users/miche/Desktop/SDM Birds/Holarctic birds 2019_08_07/Input_20190807/Mask_21.nc',
                  var="ice_thickness")

# plot
png(paste(sp_name,region,"4_plots.png",sep="_"), height=10, width=wtd,
    units='in', res=600)
par(mfrow=c(2, 2),
    oma=c(2, 0, 5, 1),
    mar=c(3, 2, 3, 3),
    mgp=c(2, 1, 0))

# occurrences
plot(WorldMap, col="cornsilk", bg="lightblue1", lwd=0.05, border="grey",
     xlim=coord[1:2], ylim=coord[3:4],
     main=paste("Clean dataset\n", dim(distrib)[1], "observations", sep=" "))
points(as.numeric(distrib[,"long"]), as.numeric(distrib[,"lat"]),
       pch="*", col=color)

# thinning
plot(WorldMap, col="cornsilk", bg="lightblue1", lwd=0.05, border="grey",
     xlim=coord[1:2], ylim=coord[3:4],
     main=paste("Thinned dataset\n", dim(ThinDistrib)[1], "observations", sep=" "))
plot(SpeciesMask, add=TRUE, col=color, border=NA)
points(as.numeric(ThinDistrib$long), as.numeric(ThinDistrib$lat),
       pch="*", col="black")

# Ensemble projection present day
plot(FlatPresRaster, main=paste("Projection present", EnsembleMethods[maxTSS], sep=" - "),
     axes=FALSE, box=FALSE, colNA="lightblue1", col=c("cornsilk",color),legend=FALSE)
plot(PresIce, add=TRUE, col=c(NA,"white"), border=NA, useRaster=F, legend=FALSE)

# Ensemble projection past
plot(FlatPastRaster, main=paste("Projection LGM", EnsembleMethods[maxTSS], sep=" - "),
     axes=FALSE, box=FALSE, colNA="lightblue1", col=c("cornsilk",color),legend=FALSE)
plot(PastIce, add=TRUE, col=c(NA,"white"), border=NA, useRaster=F, legend=FALSE)

# title
title(main=sp,cex.main= 3,outer=TRUE, line=2)
title(sub=paste("Area (square km):  LGM=",
                format(PastAreaKm, big.mark=","), "  Present = ",
                format(PresAreaKm, big.mark=","),
                "\n% of present range covered during LGM = ",
                round(overlap, digits=3), sep=""),
      outer=TRUE, line=1, cex.sub=1.5)
```

```r
  dev.off()

  #==============================#
  # 5. Plot overlap present LGM #
  #==============================#

  FlatPastRaster[is.na(FlatPastRaster)] <- 0
  FlatOverlapRaster <- (FlatPastRaster*2)+FlatPresRaster

  # plot
  png(paste(sp_name, region, "final_plot.png",sep="_"), height=6.4, width=wtd*0.75,
      units='in', res=600)

  plot(FlatOverlapRaster, col=c(cols[c(1,3,2,4)]), colNA="lightblue1",
       legend=FALSE, main=sp, box=FALSE, axes=FALSE,
       sub=paste("Area (square km):  LGM=",
             format(PastAreaKm, big.mark=","), "  Present=",
             format(PresAreaKm, big.mark=","), "\n% of present range covered during LGM = ",
             round(overlap, digits=3), sep=""))
  legend("bottomright", c("Present", "LGM", "Overlap"), fill=cols[c(3,2,4)], border=NA,
         bty="n", ncol=3)

  dev.off()

  # overwrite species file including results
  write.csv(csv, file="Species.csv", quote=FALSE, row.names=FALSE)

}
```
